# Identification of novel ubiquitin receptors on the 26S proteasome by photo-crosslinking mass spectrometry

**DOI:** 10.64898/2025.12.19.695558

**Authors:** Nicole S. MacRae, Ken C. Dong, Hiromitsu Harimoto, Andreas Martin

## Abstract

The 26S proteasome is the endpoint of the ubiquitin-proteasome system, an essential pathway for maintaining cellular homeostasis through targeted degradation of misfolded, damaged, and obsolete proteins. Substrates labeled with ubiquitin are directed to the 26S proteasome by binding to one or more ubiquitin receptors. However, ubiquitin-dependent degradation occurs even when the canonical receptor sites are mutated, suggesting the presence of additional, unidentified binding sites. Here we created photo-crosslinkable probes for ubiquitin interactions by incorporating the unnatural amino acid p-benzoyl-L-phenylalanine into ubiquitin. We show that these probes can be used to measure apparent affinities for known receptors and to reveal novel ubiquitin-binding sites on the yeast 26S proteasome. Through photo-crosslinking mass-spectrometry experiments we identified a groove on the top of the proteasome, formed by Rpn2, Rpn9, Rpn10, and Rpn12, that serves as an additional ubiquitin-binding interface. Our photo-crosslinkable probes thus serve as versatile tools for the characterization of ubiquitin-protein interactions and the identification of ubiquitin-binding domains.

## Introduction

The Ubiquitin-Proteasome system (UPS) is responsible for the degradation of damaged and obsolete proteins, and therefore plays essential roles in maintaining cellular homeostasis and the regulation of numerous vital processes ^1,2^. Dysfunction of the UPS can cause protein aggregation, neurodegenerative diseases, and various cancers. Most proteins are targeted to the 26S proteasome through the enzymatic attachment of ubiquitin to one or more of its lysine residues, with additional ubiquitins then added to lysines in ubiquitin itself to form poly-ubiquitin chains of different linkage types ^3^. The 26S proteasome consists of the 20S core particle (CP), which is a barrel shaped compartmental protease, and the 19S regulatory particle (RP), which recruits ubiquitinated substrates, deubiquitinates them, and uses ATP hydrolysis for their mechanical unfolding and translocation into the 20S CP for proteolytic cleavage ^4,5^. The 20S core consists of four stacked hetero-heptameric rings: two outer rings formed by the subunits α1-α7 (SCL1, PRE8, PRE9, PRE6, PUP2, PRE5, and PRE10 in yeast *S. cerevisiae*), and two inner rings with the subunits β1-β7 (PRE3, PUP1, PUP3, PRE1, PRE2, PRE7 and PRE4 in yeast), of which β1, β2, and β5 have proteolytic activity. The 19S RP can be further split into two subcomplexes, the base and the lid. The base contains the heterohexameric AAA+ ATPase motor formed by Rpt1-Rpt6, the large scaffolding subunit Rpn2, and two characterized ubiquitin receptors, Rpn1 and Rpn13 ^6–9^. The lid consists of eight scaffolding proteins, Rpn3, 5, 6, 7, 8, 9, 12, and Sem1, and the essential deubiquitinase Rpn11. A third, integral Ub receptor, Rpn10, bridges the base and lid in the assembled 19S RP ^10,11^. To be suited for proteasomal turnover, substrates need to contain a bipartite degradation signal consisting of poly-ubiquitin or multiple mono-ubiquitin modifications and an unstructured initiation region for engagement by the ATPase motor ^12^. Upon ubiquitin binding to one or more receptors, the initiation region enters the central channel of the ATPase hexamer, which triggers a major conformational change of the 19S RP from a resting state (s1) to the processing states (non-s1) that subsequently drive the mechanical substrate threading into the 20S CP ^5^.

Ubiquitin interactions with the three canonical receptors, Rpn10, Rpn13, and Rpn1, were previously characterized by nuclear magnetic resonance (NMR) spectroscopy, isothermal calorimetry (ITC), and surface plasmon resonance (SPR) ^6,7,13–15^, which identified specific residues responsible for their unique ubiquitin-binding behaviors. Rpn10 binds ubiquitin through residues 228-232 in its C-terminal ubiquitin-interacting motif (UIM), while using its N-terminal Von Willebrand factor, type A (VWA) domain to interact with subunits of the 19S RP ^13,15^. Rpn13 is bound to Rpn2 at the top of the proteasome and interacts with ubiquitin through the S2-S3, S4-S5, and S6-S7 loops of its Pleckstrin-like Receptor for Ubiquitin (Pru) domain ^7,14,16^. Rpn1 binds to ubiquitin and the ubiquitin-like (UBL) domain of the cofactor Rad23 through its T1 site, and utilizes its T2 site to interact with the UBL domain of the deubiquitinase Ubp6 ^6^. It has also been reported that an N-terminal fragment of Rpn1, called the N1 site (Rpn1^214–335^), serves as an additional receptor for ubiquitin and UBL domains ^17^. All three receptors thereby contact a common interface on ubiquitin, the I44 patch ^18^, yet additional interactions confer some ubiquitin-linkage specificities, with Rpn1 and Rpn10 showing higher affinities for K48-linked ubiquitin chains, whereas Rpn13 seems to be less selective for this linkage type ^19^.

Interestingly, while mutation of all three ubiquitin receptors in *S. cerevisiae* causes decreased rates of ubiquitin-dependent degradation and increased canavanine sensitivity, the triple mutant is not fatal and does not completely eliminate ubiquitin-dependent substrate degradation by the proteasome, suggesting that there are additional, unidentified or “cryptic” receptors ^6^. Although the identification and characterization of ubiquitin-binding proteins has been enabled by pulldowns, NMR, and ITC, we aimed to develop a more generic tool for probing ubiquitin receptors and identifying novel binding interfaces on ubiquitin-interacting proteins and complexes, such as the proteasome.

In order to build a probe for ubiquitin receptors, we sought to incorporate a photo-crosslinkable unnatural amino acid in key positions within ubiquitin. We chose p-benzoyl-L-phenylalanine (BPA), which reacts with carbon-hydrogen bonds and allows crosslinking to any amino acid within a ∼ 3 Å radius when excited by 365 nm light ^20,21^. BPA has previously been used to capture protein interactions ^22^, including with ubiquitin ^23–26^, and for substrate crosslinking to the proteasome ^27^. Here, we incorporated BPA at four different sites in ubiquitin to develop photo-crosslinkable probes for the identification and characterization of ubiquitin-binding domains (UBDs). Due to BPA’s rather low crosslinking efficiency, we were able to reasonably recapitulate the ubiquitin-binding affinities of canonical proteasome receptors that were previously determined by other biochemical methods, highlighting the potential of these probes for characterizing ubiquitin-UBD interactions in general. We then used these probes to identify novel ubiquitin-binding interfaces on the 26S proteasome and found a groove formed by Rpn2, Rpn9, Rpn10, and Rpn12 on top of the proteasome, which may be used for multivalent binding of longer or branched ubiquitin chains during substrate delivery.

## Results

### Design and development of photo-crosslinkable ubiquitin probes

To develop probes for ubiquitin interactions, we used genetic code expansion to incorporate the unnatural amino acid BPA into four ubiquitin constructs, which also contained an N-terminal GlyGly extension for fluorescein conjugation by a sortase reaction and a C-terminal His_6_-tag for purification and pulldowns (Figure 1A). We used the orthogonal aminoacyl-tRNA synthetase (MjTyrRS)/tRNA pair that leads to the incorporation of BPA at any position in the coding sequence where an amber stop codon (UAG) was introduced ^21,27^ and thus allows the site-specific placement of the crosslinker in ubiquitin. We designed three probes containing BPA in or near the canonical hydrophobic I44 patch on ubiquitin, Ub-(L8BPA), Ub(H68BPA), and Ub(L73BPA), as well as one probe with BPA more distant from this interface, Ub(I36BPA), to test for the degree of non-specific crosslinking. All probes also contained an N-terminal fluorescein amidite (FAM) label. As many ubiquitin receptors specifically bind chains of ubiquitin, we designed two K48-linked ubiquitin dimers (Ub_2_): Ub_2_^Prox^ had the BPA incorporated at position 68 in the proximal Ub moiety, which also carried an N-terminal FAM label and a C-terminal His_6_-tag to prevent further conjugation; Ub_2_^Dist^ contained BPA at position 73 in the distal Ub moiety, together with a K48R mutation to prevent ubiquitin additions beyond the dimer (Figure 1B, S1A). Ub_2_^Prox^ was synthesized by conjugating Ub(K48R) to FAM-Ub(H68BPA)-His_6_, while Ub_2_^Dist^ was synthesized by conjugating Ub(L73BPA, K48R) to FAM-Ub-His_6_, both using Uba1 and the K48-specific E2 enzyme Cdc34. Since the BPA-incorporation sites in Ub(L8BPA), Ub(H68BPA), and Ub(L73BPA) are close to, but not overlapping with the hydrophobic binding interface of ubiquitin, all three variants were expected to interact with Rpn1, Rpn10, and Rpn13 at the known ubiquitin-binding sites to allow for efficient crosslinking (Figures 1C-E).

**Figure 1:**
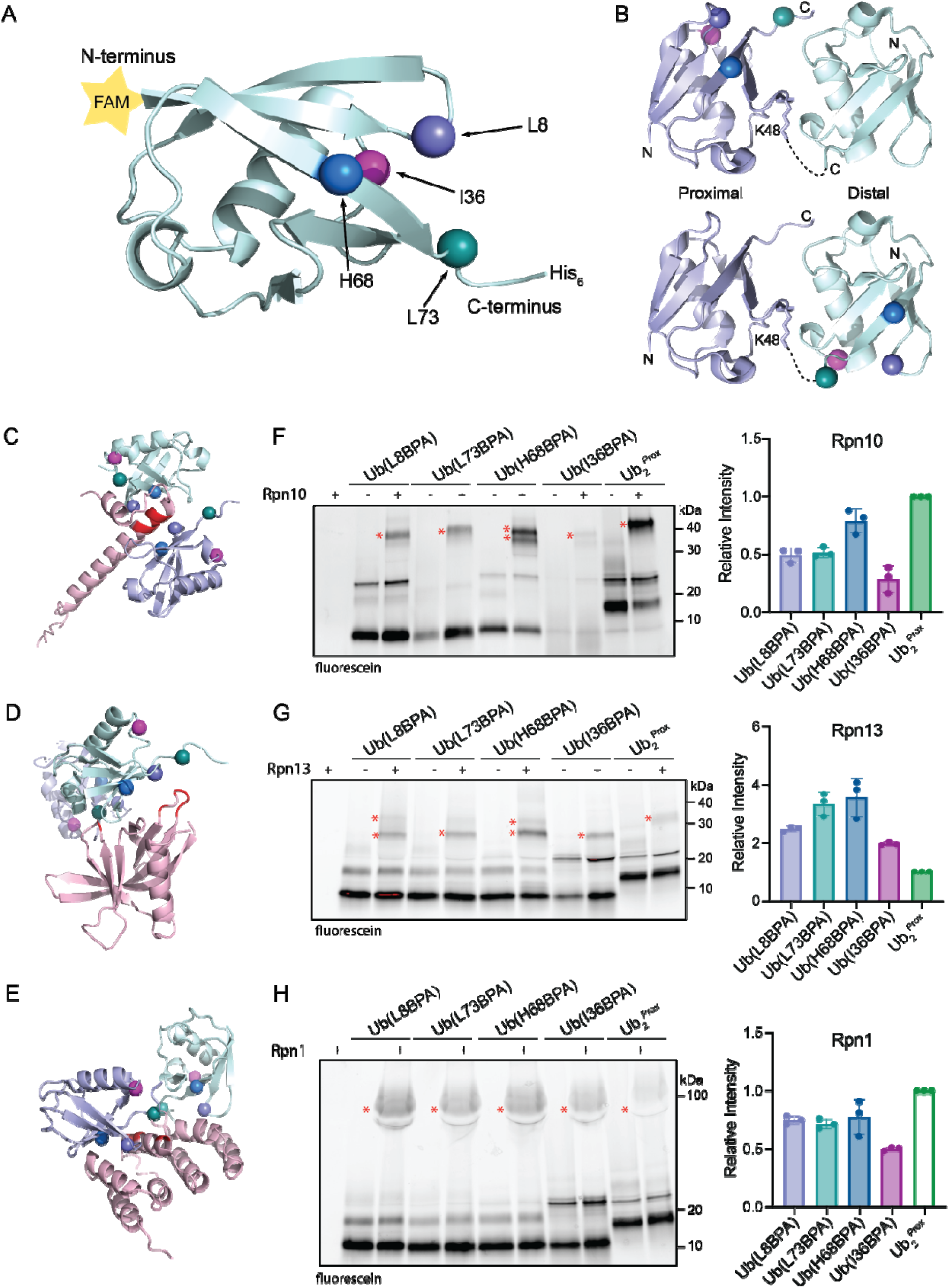
Development of photo-crosslinkable ubiquitin probes. **A)** Structure of ubiquitin (PDB: 1UBQ) with positions of BPA incorporation indicated by spheres and the N-terminal FAM label represented by a yellow star. **B)** BPA incorporation into proximal and distal ubiquitin moieties of a K48-linked di-ubiquitin. Shown is the crystal structure of di-ubiquitin (PDB: 7S6O) with the BPA locations indicated by spheres. **C)** AlphaFold model of Ub_2_ (proximal Ub in purple, distal Ub in light blue) bound to the UIM of yeast Rpn10 (pink). The residues of the UIM responsible for Ub binding (L228, A229, M230, A231, L232) are shown in red and the BPA positions in Ub_2_ are shown by spheres. **D)** Structure (PDB: 6UYI) of Ub_2_ (purple and light blue) bound to human Rpn13 (pink). The residues responsible for Ub binding (in yeast, E41, E42, L43, F45, S93) are highlighted in red. **E)** Structure (PDB: 2N3V) of Ub_2_ (purple and light blue) bound to the T1 site of yeast Rpn1 (the corresponding fragment of Rpn1 is shown in pink). Residues responsible for Ub binding (D541, D548, E552) are highlighted in red. **F)** Left: Gel analyzing the crosslinking of FAM-Ub(L8BPA), FAM-Ub(L73PA), FAM-Ub(H68BPA), FAM-Ub(I36BPA), and K48-linked FAM-Ub(H68BPA)-Ub(K48R) (Ub ^Prox^) with Rpn10. Crosslink bands are marked with *. Right: Quantification of crosslink bands, normalized to the crosslink band intensity to Ub ^Prox^. **G)** Left: Gel analyzing the crosslinking of FAM-Ub(L8BPA), FAM-Ub(L73PA), FAM-Ub(H68BPA), FAM-Ub(I36BPA), and K48-linked FAM-Ub(H68BPA)-Ub(K48R) (Ub ^Prox^) with Rpn13. Crosslink bands are marked with *. Right: Quantification of crosslink bands, normalized to the crosslink band intensity to Ub ^Prox^. **H)** Left: Gel analyzing the crosslinking of FAM-Ub(L8BPA), FAM-Ub(L73PA), FAM-Ub(H68BPA), FAM-Ub(I36BPA), and K48-linked FAM-Ub(H68BPA)-Ub(K48R) (Ub ^Prox^) with Rpn1. Crosslink bands are marked with *. Right: Quantification of crosslink bands, normalized to the crosslink band intensity to Ub ^Prox^.

Using a crosslinking assay with gel-based fluorescence readout, where a receptor is mixed with Ub(BPA), exposed to UV light (365 nm), and analyzed by SDS-PAGE, we tested how efficiently each mono-Ub probe as well as Ub_2_ ^Prox^ crosslinked to the three proteasomal receptors by tracking the FAM label on ubiquitin (Figures 1F-H, S1B-D). Ub(H68BPA) was the most efficient mono-Ub probe, and crosslinking to Rpn10 and Rpn1 could be further improved by using Ub_2_ ^Prox^. In contrast, crosslinking to Rpn13 was greatly reduced for Ub_2_ ^Prox^, likely because the Pru domain can only accommodate a single ubiquitin at a time ^14^, and the second, non-crosslinkable ubiquitin moiety in Ub_2_ ^Prox^ therefore acted as a competitor. As expected, Ub(I36BPA) with its BPA located distant from the I44 patch showed the weakest crosslinking, which was probably due to non-specific interactions at the high protein concentrations used in this assay. In summary, these initial data showed that the Ub(BPA) probes readily crosslink to known ubiquitin-binding subunits of the proteasome and can be used to capture these interactions *in vitro*.

### Ub(BPA) probes crosslink to receptors in a concentration-dependent manner

To assess whether the Ub(BPA) probes could be used to measure apparent binding affinities, we titrated the receptor subunits against a constant Ub(H68BPA) concentration in our gel-based crosslinking assay and quantified the crosslinked band to obtain a saturation curve and apparent K_D_ (K_D, app_) (Figures 2A-C). The calculated K_D, app_ values are remarkably similar to those previously determined by NMR, ITC, or SPR (Table 1) ^6,19,28,29^. Due to the rather inefficient reaction of BPA, crosslinking is slow compared to the rates for ubiquitin binding and dissociation, and the extent of crosslinking is therefore directly modulated by the K_D_.

**Figure 2:**
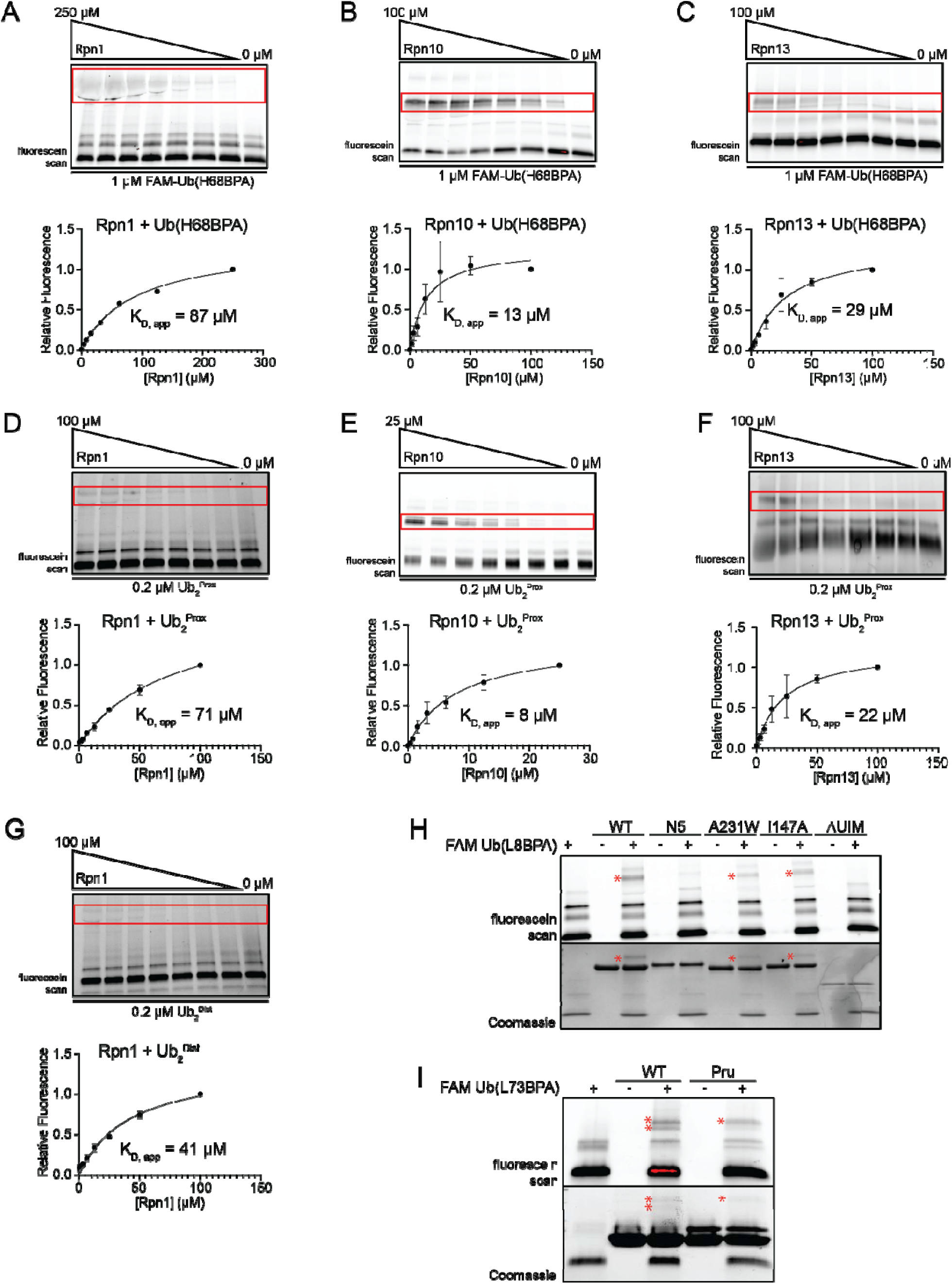
Ub(BPA) probes crosslink at the expected affinities. **A)** Top: Representative gel of a titration of Rpn1 crosslinked to FAM-Ub(H68BPA). Crosslink band is boxed in red. Bottom: Relative band intensities plotted against Rpn1 concentration and fit to a binding curve to determine K_D, app_. Shown are the average for n = 3 technical replicates. **B)** Top: Representative gel of a titration of Rpn10 crosslinked to FAM-Ub(H68BPA). Crosslink band is boxed in red. Bottom: Relative band intensities plotted against Rpn10 concentration and fit to a binding curve to determine K_D, app_. Shown are the average for n = 3 technical replicates. **C)** Top: Representative gel of a titration of Rpn13 crosslinked to FAM-Ub(H68BPA). Crosslink band is boxed in red. Bottom: Relative band intensities plotted against Rpn13 concentration and fit to a binding curve to determine K_D, app_. Shown are the average for n = 3 technical replicates. **D)** Top: Representative gel of a titration of Rpn1 crosslinked to Ub ^Prox^. Crosslink band is boxed in red. Bottom: Relative band intensities plotted against Rpn1 concentration and fit to a binding curve to determine K_D, app_. Shown are the average for n = 3 technical replicates. **E)** Top: Representative gel of a titration of Rpn10 crosslinked to Ub ^Prox^. Crosslink band is boxed in red. Bottom: Relative band intensities plotted against Rpn10 concentration and fit to a binding curve to determine K_D, app_. Shown are the average for n = 4 technical replicates. **F)** Top: Representative gel of a titration of Rpn13 crosslinked to Ub ^Prox^. Crosslink band is boxed in red. Bottom: Relative band intensities plotted against Rpn13 concentration and fit to a binding curve to determine K_D, app_. Shown are the average for n = 3 technical replicates. **G)** Top: Representative gel of a titration of Rpn1 crosslinked to Ub ^Dist^. Crosslink band is boxed in red. Bottom: Relative band intensities plotted against Rpn1 concentration and fit to a binding curve to determine K_D, app_. Shown are the average for n = 3 technical replicates. **H)** Crosslinking of FAM-Ub(L8BPA) to WT Rpn10 and its N5, A231W, I147A, and ΔUIM mutants. Crosslink bands are indicated with *. **I)** Crosslinking of FAM-Ub(L8BPA) to WT Rpn13 and its Pru-domain mutant. Crosslink bands are indicated with *.

**Table 1:** Calculated K_D, app_ values compared to previously established K_D_ values.

| | Apparent $K_D$ ($K_{D, app}$) | Published $K_D$ |
| --- | --- | --- |
| Rpn10 – Ub(H68BPA) | $13 \pm 7 \mu\text{M}$ | $45 \mu\text{M}$ |
| Rpn10 – Ub<sub>2</sub><sup>Prox</sup> | $8 \pm 6 \mu\text{M}$ | $13 \mu\text{M}$ |
| Rpn13 – Ub(H68BPA) | $29 \pm 8 \mu\text{M}$ | $91 \mu\text{M}$ |
| Rpn13 – Ub<sub>2</sub><sup>Prox</sup> | $22 \pm 8 \mu\text{M}$ | $29 \mu\text{M}$ |
| Rpn1 – Ub(H68BPA) | $87 \pm 20 \mu\text{M}$ | $120 \mu\text{M}$ |
| Rpn1 – Ub<sub>2</sub><sup>Prox</sup> | $71 \pm 13 \mu\text{M}$ | $12 \mu\text{M}$ |
| Rpn1 – Ub<sub>2</sub><sup>Dist</sup> | $41 \pm 10 \mu\text{M}$ | $12 \mu\text{M}$ |

While the K_D, app_ for monomeric Ub(H68BPA) crosslinking to Rpn1 was less than 1.5-fold below the published K_D_, the K_D, app_ for Rpn10 and Rpn13 showed slightly larger deviations, ∼ 3 - 3.5-fold lower than previously reported values, which is likely due to the irreversible nature of the crosslink. As expected, a lower K_D, app_ was observed for the dimeric Ub_2_^Prox^ binding to Rpn10, in good agreement with the published data. For the interaction with Rpn13, Ub_2_^Prox^ showed a lower value than Ub(H68BPA), which is consistent with a potential competition between the BPA-containing and BPA-free ubiquitin moieties for Pru-domain binding. For crosslinking of the ubiquitin dimer to Rpn1 we observed a clear positional difference, where Ub_2_^Dist^ had an almost 2-fold lower K_D, app_ than Ub_2_ ^Prox^ (Figures 2D, G; Table 1). This agrees with the structure of K48-linked Ub_2_ bound to the T1 site of Rpn1 (Figure 1E), showing that it is the distal moiety that primarily contacts the canonical ubiquitin-binding residues of the receptor.

Overall, our measured affinities indicate that fluorescence-based photo-crosslinking of the Ub(BPA) probes can serve as a valuable orthogonal method for the experimental determination of approximate binding affinities.

### Ub(BPA) crosslinking relies on canonical Ub-receptor interactions

To further assess whether Ub(BPA) crosslinking is dependent on specific ubiquitin-UBD interactions, we mutated residues in Rpn10 and Rpn13 that were previously shown to be responsible for binding ^6,7,13,14^. Ubiquitin interactions with Rpn10 had been mapped to five amino acids in the UIM ^29^, and our mutation of all five residues (L228N, A229N, M230N, A231N, L232N – labeled “Rpn10 N5”) indeed completely abolished observable crosslinking (Figure 2H). Mutation of just one of these residues, A231W, led to a considerable decrease in the crosslinking intensity. In contrast, mutation of I147, which is located in the VWA domain of Rpn10 and was previously proposed as part of additional ubiquitin-binding interface ^30^, had no major effect on crosslinking, suggesting that the VWA domain is not involved in ubiquitin binding for substrate recruitment to the proteasome. Consistently, Ub(BPA) showed no detectable crosslinking to the isolated VWA domain (ΔUIM) (Figure 2H).

For Rpn13, we mutated five Pru-domain residues involved in ubiquitin binding (E41K, E42K, L43A, F45A, S93D) and observed a significant reduction in crosslinking compared to the wild-type Rpn13 (Figure 2I). In addition to the mutation-sensitive crosslink band, Rpn13 exhibited a secondary band at higher molecular weight that we attribute to non-specific crosslinking, either due to the high protein concentrations used in our assay or the exposed interfaces on isolated Rpn13 that are otherwise used for receptor interactions with the proteasome.

### LC-MS/MS can be used to identify ubiquitin-crosslinked peptides in Rpn10

Rpn10 is the best-characterized ubiquitin receptor and thus can serve as a model system for testing whether Ub(BPA) crosslinking can be combined with mass spectrometry to identify specific binding sites on the proteasome. We therefore crosslinked Ub(L73BPA) to Rpn10, digested the reaction products with chymotrypsin, ran LC-MS/MS analyses, and used the pLink software ^31^ to assign specific spectra to crosslinked peptides. This analysis identified seven Rpn10 peptides, whose mapping on the AlphaFold model of the ubiquitin-bound Rpn10 structure revealed that three of them lie in or adjacent to the UIM (P222, E227, R247), while the other four peptides stem from crosslinks to the VWA domain (A5, V49, A94, K104; Figure 3A). The crosslinks in and near the UIM confirm the UIM as the ubiquitin-binding site (Figures 3B-D). That the identified residues do not agree with those previously shown to directly contact ubiquitin is expected, given our experimental design, with BPA positions on ubiquitin bordering yet not overlapping with the interaction interface. Ub(BPA) crosslinking can thus detect broad regions of interaction, but not pinpoint specific residues. The observed crosslinks to the VWA domain are likely non-specific and caused by Rpn10 being outside the proteasome context, as these positions are buried when Rpn10 is bound to the 19S RP. Due to its high sensitivity, mass spectrometry detected those off-target crosslinks, whereas no significant crosslinking was observed for the Rpn10 ΔUIM construct in our gel-based assay (Figure 2H).

**Figure 3:**
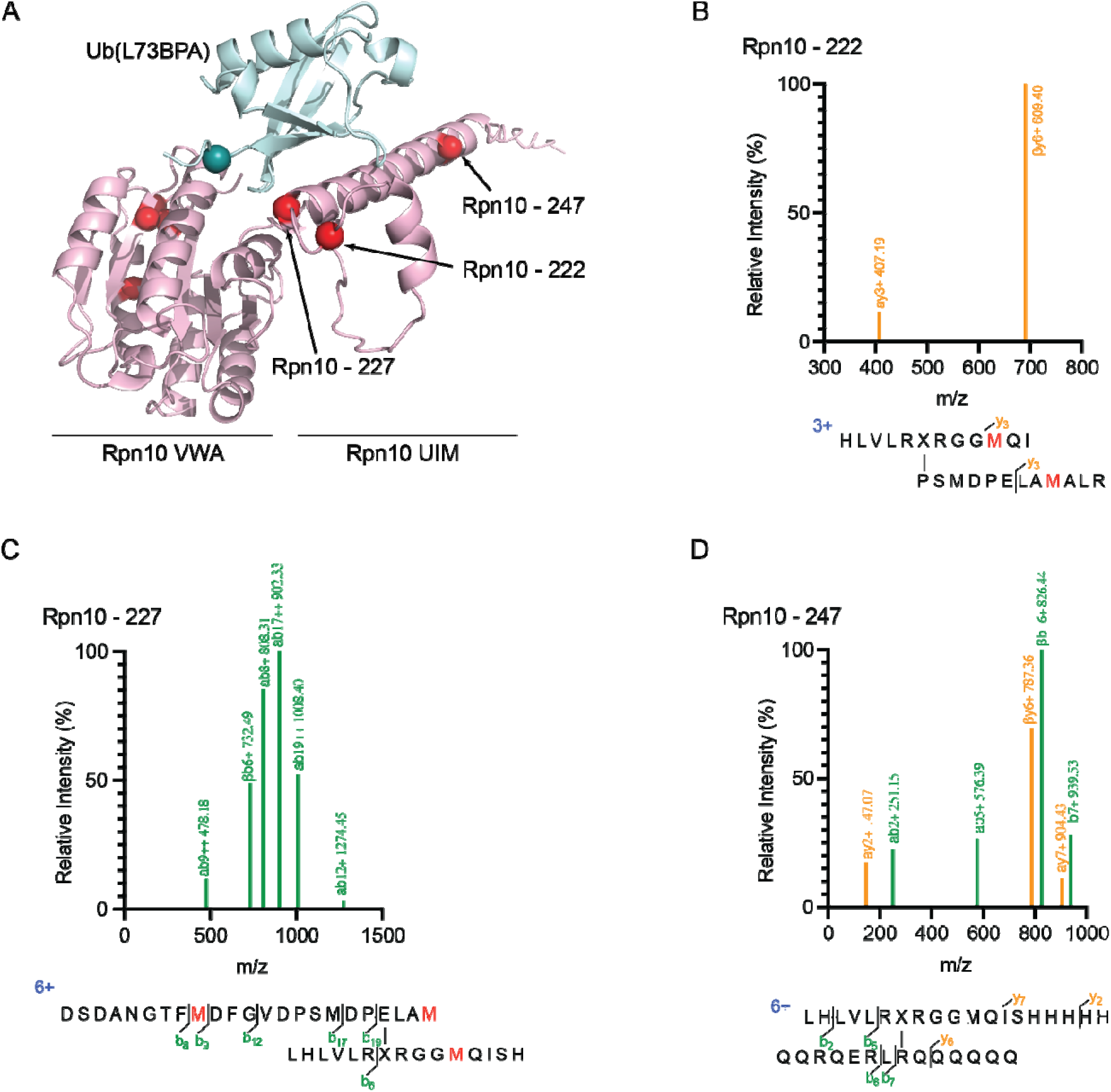
Identification of crosslinked peptides from Rpn10 using LC-MS/MS. **A)** AlphaFold structural model of Rpn10 (pink) bound to Ub (light blue). L73BPA in ubiquitin is shown as a teal sphere and identified crosslinks in Rpn10 are shown as red spheres. **B)** Left: Spectrum of an identified peptide containing a crosslink at Rpn10 222. Right: Schematic of the identified peptide. BPA is indicated as ‘X’. **C)** Left: Spectrum of an identified peptide containing a crosslink at Rpn10 227. Right: Schematic of identified peptide. BPA is indicated as ‘X’. **D)** Left: Spectrum of an identified peptide containing a crosslink at Rpn10 247. Right: Schematic of identified peptide. BPA is indicated as ‘X’.

### Ub(BPA) probes identify novel Ub receptors on the 26S proteasome

To reveal novel ubiquitin receptors and binding sites on the 26S proteasome, we crosslinked Ub(L73BPA) to endogenous proteasome holoenzymes from yeast. These proteasomes were purified by affinity chromatography, using a FLAG-tag on Rpn11, and size-exclusion chromatography (Figure S3A, S3B), such that all potentially associated proteins, including the shuttle factors Dsk2 and Rad23, the deubiquitinase Ubp6, or the E3 ligase Hul5, were maintained. Exposing these proteasomes in ATP to UV-induced crosslinking with FAM-Ub(L73BPA) for 30 minutes produced a continuous smear on the SDS-PAGE gel up to higher molecular weights, indicating that the probe reacted with multiple components of the proteasome (Figure 4A). We also found that the proteasome crosslinking pattern on the gel changed dramatically in the presence of non-hydrolyzable ATPγS (Figure 4B), which is known to shift the conformational equilibrium from the resting (s1) state to a processing (s4) state ^32,33^. These changes suggest a difference in the availability of Ub-binding sites between resting and processing states, and support a model where the conformational transitions to non-s1 states after substrate insertion into the ATPase motor facilitate ubiquitin release from certain receptors or interfaces ^34^. In order to identify which subunits were crosslinked to our ubiquitin probe, we developed a workflow for sample preparation that capitalized on the His_6_-tag present at the C-terminus of the probe (Figure S3C). Purified proteasomes were crosslinked to Ub(L73BPA) under the standard conditions used in our other assays and then completely denatured with 6 M guanidinium chloride. The Ub probe and any crosslinked proteins were enriched by Ni-NTA affinity purification, trypsinized, and analyzed by LC-MS/MS. The pLink search for crosslinked peptides varied between individual attempt, and each proteasomal subunit was therefore searched four times. A particular peptide was only considered a positive hit if it was present in at least three of the four independent searches.

**Figure 4:**
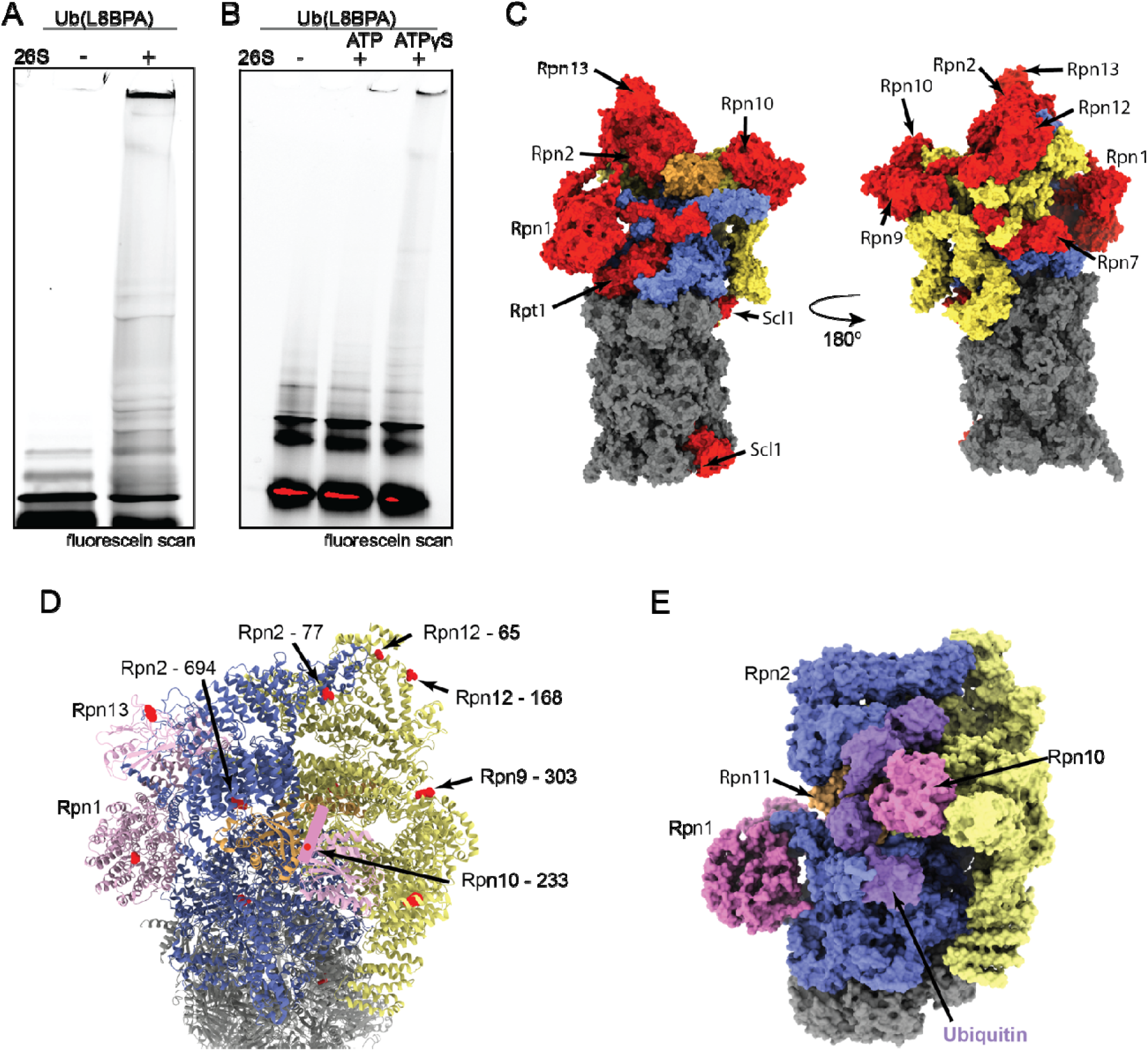
Ub(BPA) probes identify novel ubiquitin receptors on the 26S proteasome. **A)** Gel of Ub(L8BPA) crosslinked to purified 26S proteasome imaged with fluorescein fluorescence scan. **B)** Crosslinking of Ub(L73BPA) to purified 26S proteasome in the presence of ATP (stabilizing primarily the resting s1 state) or in the presence of ATPγS (stabilizing the processing s4 state) imaged by fluorescein fluorescence scan. **C)** Front view (left) and back view (right) of the 26S proteasome structure (PDB: 6FVT) with crosslinked subunits shown in red, Rpn11 in orange, non-crosslinked base subunits in blue, non-crosslinked lid subunits in yellow, and non-crosslinked core subunits in grey. **D)** View from the top right of the front orientation shown in (C), with all subunits shown in ribbon representations and colored as in (C), and with identified crosslinks indicated by red spheres. Rpn10’s UIM (unresolved in this structure, PDB: 6FVT) is depicted as a pink cylinder. Crosslinks surrounding the groove atop the regulatory particle are labeled. **E)** Structure of the human 26S proteasome with a bound K11/K48-linked branched ubiquitin chain (PDB: 8JTI). Ubiquitin moieties are shown in purple, Rpn1 and Rpn10 in pink, other base subunits in blue, Rpn11 in orange, other lid subunits in yellow, and core subunits in grey.

Using this workflow, we identified crosslinked peptides from numerous subunits of the 19S regulatory particle, including the canonical Ub receptors Rpn1, Rpn10, and Rpn13, the base subunits Rpn2 and Rpt1, the lid subunits Rpn7, Rpn9, and Rpn12, and the α1 subunit of the 20S core (Figure 4C; Table 2; Figure S6A-I). Rpn10 had a crosslink at A233, in the middle of its UIM (Figure S4A), while Rpn13 showed a crosslink at E60, located adjacent to the Pru residues (Figure S4B). The crosslinked residue in Rpn1, L873, lies in a cleft between Rpn1’s toroid and N-terminal domains, near but not immediately adjacent to the T1 site (Figure S4C). Given this position of L873, we assume that other, more exposed residues may have been responsible for the robust Ub crosslinking observed for Rpn1 in our gel-based assays, yet were not identified due to the intricacy of detecting crosslinked peptides by mass spec. Additionally, we identified peptides from Ubp6 and Hul5 (Table 2). The crosslink on Ubp6 is located adjacent to its ubiquitin-binding site (Figure S5A), whereas the crosslink on Hul5 lies in a secondary site predicted by AlphaFold2 to interact with the E2 enzyme Ubc4 (Figure S5B).

**Table 2:**
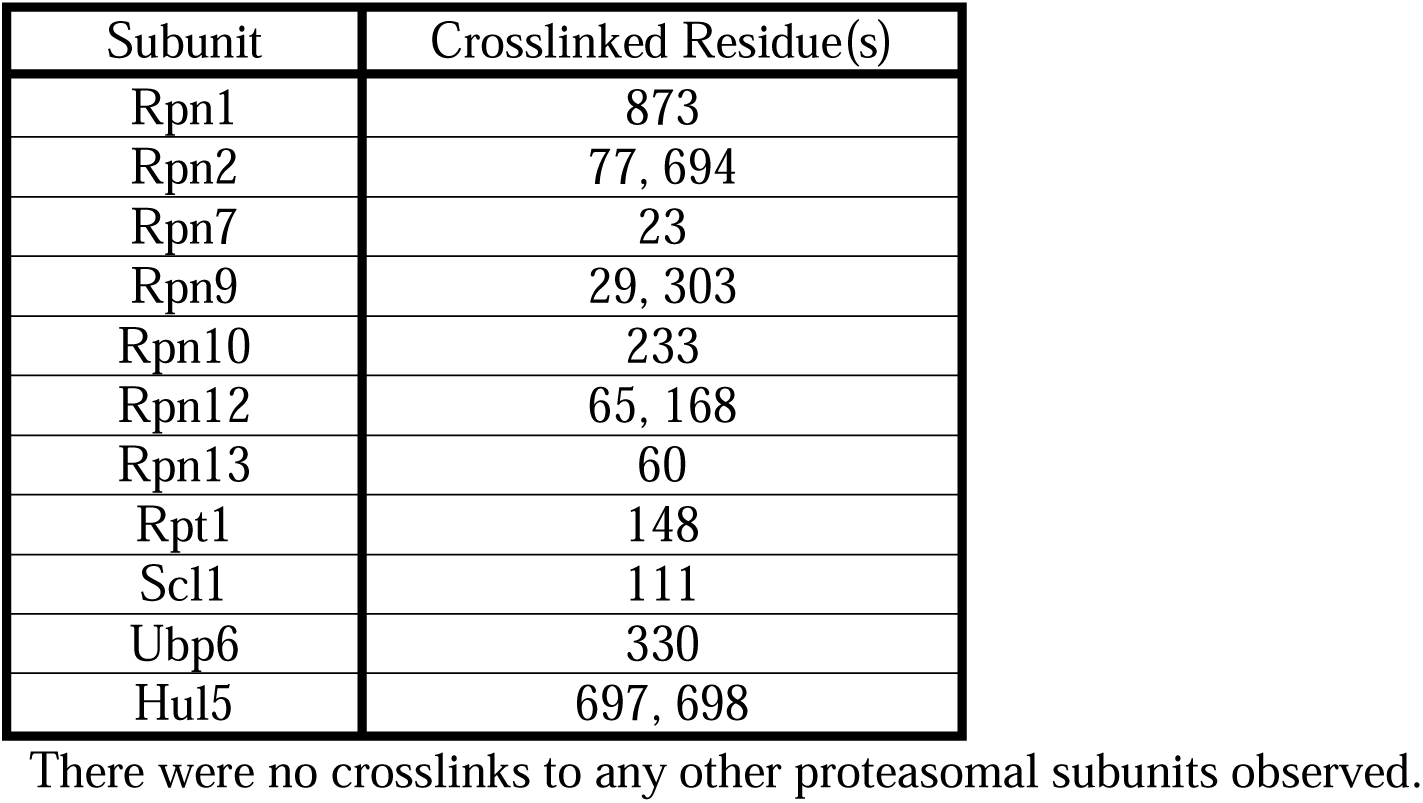
Crosslinks identified from pLink searches.

Mapping these crosslinks onto the structure of the 26S proteasome in the resting s1 state revealed that several contact points surround a groove formed by Rpn2, Rpn9, Rpn10, and Rpn12 on top of the 19S RP (Figure 4D). This groove may serve as a binding region for extended or branched ubiquitin chains that interact with Rpn10’s UIM, which lies just below the groove. Indeed, recent cryo-EM structures of the human 26S proteasome with branched K11/K48-linked chains show ubiquitin moieties positioned along the same groove ^35,36^ (Figure 4E), in nice agreement with our crosslinking data. Additional crosslinks we observed in the back of this groove suggest that ubiquitin chains longer than the ones used in these structural studies may extend further toward the back of the proteasome. Together with the structural data, our crosslinking results suggest that the groove flanked by Rpn2, Rpn9, Rpn10, and Rpn12 serves as a novel ubiquitin-binding region on top of the 26S proteasome. Moreover, these findings further support the potential of Ub(BPA) crosslinking for mapping ubiquitin-interacting sites on macromolecular complexes.

## Discussion

Due to ubiquitin’s promiscuity and often low binding affinities, its interactions with other proteins have been difficult to capture and characterize biochemically. The use of traditional chemical crosslinkers to capture non-enzymatic ubiquitin interactions is limited by large crosslinking radii and relies on the presence of specific functional groups to crosslink between. Our results show that BPA incorporation near the canonical interface of ubiquitin allows the development of specific interaction probes. Ub(BPA) crosslinking can thereby be used not only to identify novel binding sites, but also to estimate binding affinities with a gel-based fluorescence assay that provides K_D_ values similar to those from conventional methods, like NMR, ITC, or SPR, which require specialized equipment and expertise.

Our assay is not an equilibrium method due to the irreversibility of crosslinking. However, because of the slow reaction of BPA, the extent of crosslinking is modulated by the pre-equilibrium of ubiquitin binding and dissociation, such that the derived K_D, app_ values are in reasonable agreement with the binding affinities determined by true equilibrium measurements. Furthermore, our data show that positional incorporation of Ub(BPA) into a dimer revealed different K_D, app_ values, providing a platform for determining the binding contributions of particular ubiquitin moieties within a chain.

BPA is more specific than traditional chemical crosslinkers due to its short crosslinking radius, yet retains promiscuity by reacting indiscriminately with C-H bonds. BPA crosslinking is therefore an effective approach for capturing low- to medium-affinity interactions, such as ubiquitin binding. When we applied Ub(BPA) crosslinking to the 26S proteasome, we detected all three Ub receptors, indicating that these BPA probes nicely complement conventional chemical crosslinkers ^37^ and alternative crosslinkable ubiquitin variants ^38^. We also identified crosslinks to several other proteasome-resident proteins that are known to interact with ubiquitin, such as Ubp6 and Hul5, highlighting how Ub(BPA) can capture small populations of canonical ubiquitin-protein interactions *in vitro*. Moreover, our crosslinked peptide search identified additional binding sites, suggesting that a groove on the top of the regulatory particle interacts with ubiquitin. Two recent structures showed that K11/K48-branched Ub chains bind to the human 26S proteasome through the front of this groove ^35,36^. These structures thus provide some validation of our crosslinking / mass-spectrometry hits and indicate that this is a conserved ubiquitin-binding groove shared between human and yeast proteasomes. It is possible that this groove accommodates chains with specific linkages or topologies to select certain substrates for degradation.

Future work will have to investigate the context of ubiquitin binding to the groove on top of the 26S proteasome. Additionally, our Ub(BPA) probes can be used as a starting point to synthesize more complex probes with Ub(BPA) incorporated into Ub chains that contain certain physiologically relevant features, including branches. These more complex probes and the Ub(BPA) presented here could be applied to other systems for mapping ubiquitin-binding sites and determining which moieties in chains are contacted. However, it is important to note that several of our identified crosslinks, especially for insolated receptor subunits, are unlikely to be physiologically relevant, given their expected steric clashes with other subunits in the context of the 26S proteasome holoenzyme. This highlights how photo-crosslinking/mass spectrometry can capture off-target false positives and that it is necessary to validate identified crosslinks through additional mutational, biochemical, or structural studies when applying these probes to map novel ubiquitin interactions.

## Methods

### Expression and purification of Ub(BPA)

BL21 DE3* competent cells were co-transformed with vectors coding for the BPA aminoacyl-tRNA synthetase (MjTyrRS)/tRNA pair and ubiquitin containing an amber stop codon in the position desired for BPA incorporation, Ub(BPA) ^21,27^. Cells were grown in dYT media at 37 °C until OD_600_ ∼0.6, then pBPA dissolved in NaOH was added to a final concentration of 2 mM, and cells were induced with 1 mM final IPTG overnight at 18 °C. Cells were harvested by centrifugation and resuspended in NiA buffer (50 mM HEPES pH 7.6, 250 mM NaCl, 20 mM imidazole) with protease inhibitors (AEBSF, leupeptin, pepstatin, aprotinin). Pellets were lysed by sonication, and the lysate was clarified by centrifugation. His-tagged Ub(BPA) proteins were affinity purified using Ni-NTA resin, washed with NiA buffer and eluted with NiB buffer (50 mM HEPES pH 7.6, 250 mM NaCl, 250 mM imidazole), followed by size-exclusion chromatography using a Superdex 75 16/600 column (Cytiva) equilibrated in GF buffer (50 mM HEPES pH 7.6, 50 mM NaCl, 80 mM KCl, 10 mM MgCl_2_, 5% glycerol). Fractions were concentrated using a 3K cutoff concentrator. Ub(BPA) variants without a His_6_ tag were purified by acetic acid precipitation, followed by overnight dialysis in Buffer A1 (50 mM NaOAc), ion-exchange chromatography using a 5 mL HiTrap SP FF with Buffer A1 and 0-70% Buffer B1 (50 mM NaOAc, 500 mM NaCl) over 20 column volumes, and size-exclusion chromatography using a Superdex 75 16/600 column (Cytiva) equilibrated in GF buffer. Fractions were concentrated using a 3K cutoff concentrator.

### Fluorescent labeling of Ub(BPA)

Purified Ub(BPA) was mixed with 20 μM sortase, 500 μM fluorescein-LPETGG, and 10 mM CaCl_2_, and incubated at room temperature for 1 h. The sample was then purified by Ni-NTA affinity using a 1 mL HisTrap HP column (Cytiva) equilibrated with NiA buffer, and eluted using NiB buffer. It was further purified by size-exclusion chromatography using a Superdex S75 Increase column (Cytiva) equilibrated in GF buffer.

### Synthesis of K48-linked Di-Ub Chains

K48-linked Ub_2_ probes were synthesized by incubating 500 μM Ub(K48R) and 10 μM FAM-Ub(BPA)-His_6_ (for Ub_2_ ^Prox^) or 500 μM Ub(BPA, K48R) and 10 μM FAM-Ub-His_6_ (for Ub_2_ ^Dist^) with 0.3 μM mouse E1, 3 μM Cdc34, and 1 mM ATP in a total volume of 200 μL at 37 °C overnight ^39^. The Ub_2_ probes were cleaned up by Ni-NTA affinity purification using a 1 mL HisTrap HP column (Cytiva), followed by size-exclusion chromatography using a Superdex 75 Increase 10/300 (Cytiva) equilibrated in GF buffer. Fractions containing Ub_2_ probe were collected and concentrated using a 3K cutoff concentrator.

### Expression and Purification of Proteasome Subunits

BL21 DE3* competent cells were transformed with vectors containing His-tagged Rpn1, Rpn10, or Rpn13. Cells were grown in dYT media to OD_600_ ∼0.6 at 37°C, then induced with 1 mM IPTG, and incubated overnight at 18 °C. Cells were harvested by centrifugation and resuspended in NiA buffer (50 mM HEPES pH 7.6, 250 mM NaCl, 20 mM imidazole) with protease inhibitors (AEBSF, leupeptin, pepstatin, aprotinin). Pellets were lysed by sonication and lysate was clarified by centrifugation. His-tagged proteasome subunits were purified by affinity purification using Ni-NTA resin washed with NiA buffer and eluted with NiB buffer (50 mM HEPES pH 7.6, 250 mM NaCl, 250 mM imidazole), followed by size-exclusion chromatography using a Superdex 75 16/600 column (Rpn10, Rpn13) or a Superdex 200 16/600 (Rpn1) equilibrated in GF buffer (50 mM HEPES pH 7.6, 50 mM NaCl, 80 mM KCl, 10 mM MgCl_2_, 5% glycerol). Fractions were concentrated and concentration was determined by absorbance A_280_ measured on a Nanodrop.

### Expression and Purification of Yeast Proteasome

Yeast 26S proteasome was purified from the YYS40 strain containing FLAG-Rpn11 ^40^. Yeast cells were streaked from a glycerol stock on a YPD plate and incubated at 30 °C until colonies appeared. 5 mL starter cultures were inoculated in YPD using one colony per starter and incubated at 30 °C overnight. 50 mL starter cultures with YPD media were inoculated with the 5 mL starter cultures and incubated at 30 °C overnight. 1L YPD media was inoculated with 25 mL starter culture and incubated at 30 °C for 3 days. Cells were collected by centrifugation, resuspended in buffer, and popcorned into liquid nitrogen. After cryogrinding of the cells, the lysate powder was thawed in lysis buffer (60 mM HEPES pH 7.6, 25 mM NaCl, 25 mM KCl, 5 mM MgCl_2_, 0.5 mM EDTA, 5% glycerol, supplemented with 0.01% NP-40, 0.05 mg/ml creatine phosphokinase, 16 mM creatine phosphate, 2 mM ATP). Lysate was clarified at 15,000 rpm for 30 mins, the supernatant was incubated with FLAG resin equilibrated with yeast GF buffer (60 mM HEPES pH 7.6, 25 mM NaCl, 25 mM KCl, 5 mM MgCl_2_, 0.5 mM EDTA, 5% glycerol, 0.5 mM ATP) at 4 °C for 30 minutes. The resin was washed with yeast GF four times, the proteasome was eluted using yeast GF buffer supplemented with 0.15 mg/mL FLAG peptide, and the solution was concentrated in a 10K cutoff concentrator to a volume < 500 μL. Proteasome was cleaned up by size-exclusion chromatography using a Superose 6 Increase 10/300 column (Cytiva) equilibrated with FPLC buffer (25 mM HEPES pH 7.6, 25 mM NaCl, 25 mM KCl, 5 mM MgCl_2_, 0.5 mM EDTA, 5% glycerol, 0.5 mM ATP, 1 mM DTT) and concentrated to 100 μL using a 10K cutoff concentrator. Concentration was determined by Bradford.

### Gel-Based Crosslinking Assay

Purified Ub(BPA) probe and proteasome subunits or 26S proteasome holoenzyme were diluted to indicated concentrations in GF buffer in a total volume of 10 μL. Reactions were exposed to UV light (365 nm) on ice for 30 minutes. Crosslinking was analyzed by SDS-PAGE and imaging of the gels by fluorescein fluorescence, before staining with Coomassie. Band intensities on gels were quantified with BioRad Image Lab and GraphPad Prism.

### Rpn10 Crosslinking

80 μL of 3.25 μM Rpn10 was crosslinked to 2.5 μM Ub(L73BPA) by incubating on ice under UV light (365 nm) for 30 minutes. The sample was buffer-exchanged into 50 mM ammonium bicarbonate pH 8 using a 3K cutoff concentrator and the volume was reduced to 50 μL.

### 26S Crosslinking and Denaturing Enrichment

1.5 μM yeast 26S proteasome was crosslinked to 1 μM Ub(L73BPA) in 40 μL by incubation on ice under UV light (365 nm) for 30 minutes. The sample was diluted to 1 mL in Denaturing Buffer (50 mM HEPES pH 7.6, 20 mM imidazole, 6 M guanidine hydrochloride) and incubated at room temperature for 15 minutes. The sample was loaded onto a 1 mL HiTrap column (Cytiva) equilibrated with Denaturing Buffer, washed twice with 5 mL Denaturing Buffer, and eluted with 7 mL of 50 mM HEPES pH 7.6, 250 mM imidazole, 6 M guanidine hydrochloride. The sample was concentrated in a 3K cutoff concentrator and buffer exchanged into 50 mM ammonium bicarbonate pH 8, before reducing the volume to 50 μL.

### Tryptic Digestion

The sample volume was brought to 60 μL with 50 mM ammonium bicarbonate pH 8 and 25% acetonitrile. TCEP was added to a final concentration of 5 mM, and the sample was incubated at room temperature for 20 minutes. Iodoacetamide was added to a final concentration of 10 mM, and the sample was incubated at room temperature in the dark for 20 minutes. CaCl_2_ was added to a final concentration of 1 mM, and the sample was incubated overnight at 37 °C with at least 1 μg trypsin. The sample was acidified with formic acid to 5%, diluted, and reduced in volume to 10 μL in a Speedvac.

### High pH Offline Fractionation

In-gel trypsin-digested peptides from crosslinked 26S proteasome samples were filtered with a 0.22 μm centrifugal filter (Millipore UFC30GVNB) and manually loaded on an in-house prepared fused silica capillary column (250 μm X 35cm) with Kasil outlet end. The column was packed with ReproSil pHoenix C18-3.0 μm high PH resin (Dr. Maisch GmbH). Peptides were eluted at a flow rate of 2 μL/min using a linear gradient of 2–40% buffer B in 140 min (buffer A: 0.1% triethylamine (TEA) pH 10.0, 5% acetonitrile in water; buffer B: 0.1% TEA pH 10.0, 95% acetonitrile in water) with an Agilent 1200 nanoflow LC. Samples were collected every 3 minutes during the gradient using a fraction collector (Biorad 2110). The 60 fractions were concatenated to 8 fractions and acidified with formic acid to pH 2.0, before reducing their volume to 10 μL in a Speedvac.

### LC-MS/MS

Trypsin-digested peptides were analyzed by online capillary nanoLC-MS/MS using a 25 cm reversed phase column and a 10 cm pre-column fabricated in-house (75 µm inner diameter, packed with ReproSil-Gold C18-1.9 μm resin (Dr. Maisch GmbH)) that was equipped with a laser-pulled nano-electrospray emitter tip. The pre-column used 3.0 μm packing (Dr. Maisch GmbH). Peptides were eluted at a flow rate of 300 nL/min using a linear gradient of 2–40% buffer B in 140 min (buffer A: 0.05% formic acid, 5% acetonitrile in water; buffer B: 0.05% formic acid, 95% acetonitrile in water) in an Thermo Fisher Easy-nLC1200 nanoLC system. Peptides were ionized using a FLEX ion source (Thermo Fisher Scientific) using electrospray ionization into an Fusion Lumos Tribrid Orbitrap Mass Spectrometer (Thermo Fisher Scientific). Data were acquired in Orbi-trap mode. Instrument method parameters were as follows: MS1 resolution, 120,000 at 200 m/z; scan range, 350−1600 m/z. The top 20 most-abundant ions were subjected to collision-induced dissociation with a normalized collision energy of 35%, activation q 0.25, and precursor isolation width 2 m/z. Dynamic exclusion was enabled with a repeat count of 1, a repeat duration of 30 seconds, and an exclusion duration of 20 seconds.

### Crosslinked Peptide Search

Crosslinked peptides were identified using the pLink software ^31^. MGF files exported from PEAKS (Bioinformatics Solutions Inc.) were searched with the following parameters: crosslinker set as BPA, enzyme type was set to nonspecific, up to 3 missed cleavages, peptide mass 600 - 10000 Da, peptide length 4 – 100 residues, precursor tolerance set to 10 ppm, fragment tolerance set to 10 ppm, fixed carbaminomethyl C modification, variable oxidation M modification, with a reference database containing the Ub(BPA) sequence and the sequence of 1-2 proteasome subunits. Each subunit was included in 4 different reference databases, once as the only sequence other than Ub(BPA). Crosslinked peptides that were identified in 75-100% of searches for that subunit were considered true positive hits. Crosslinked peptides identified in 50% or fewer of searches were considered false positives. Spectra corresponding to the crosslinked peptides were analyzed in pLabel and plotted in GraphPad Prism.

## Resource availability

### Lead contact

Further information and requests for resources and reagents should be directed to and will be fulfilled by the lead contact, Andreas Martin.

### Materials and data availability

Data generated or analyzed during this study are included in this manuscript and the Supplementary materials. This paper does not report original code. All constructs are available from the lead contact upon request and completion of a Material Transfer Agreement.

## Acknowledgments

The authors thank all members of the Martin Lab for valuable discussions and Zaw Htet for creating the UIM mutations in Rpn10.

## Author Contributions

N.S.M., K.C.D., and A.M. conceptualized the study. N.S.M., K.C.D., and H.H. designed and cloned constructs. N.S.M., K.C.D., and H.H. purified protein. N.S.M. performed crosslinking assays and prepared samples for mass spectrometry. N.S.M. and K.C.D. analyzed biochemical data. N.S.M. analyzed crosslinking data and performed crosslinked peptide searches. N.S.M., K.C.D., H.H., and A.M. provided resources. A.M. supervised the study and acquired funding.

N.S.M. wrote the manuscript. N.S.M., K.C.D., and A.M. edited the manuscript and contributed to revisions.

## Supplemental Figures

**Supplemental Figure 1:**
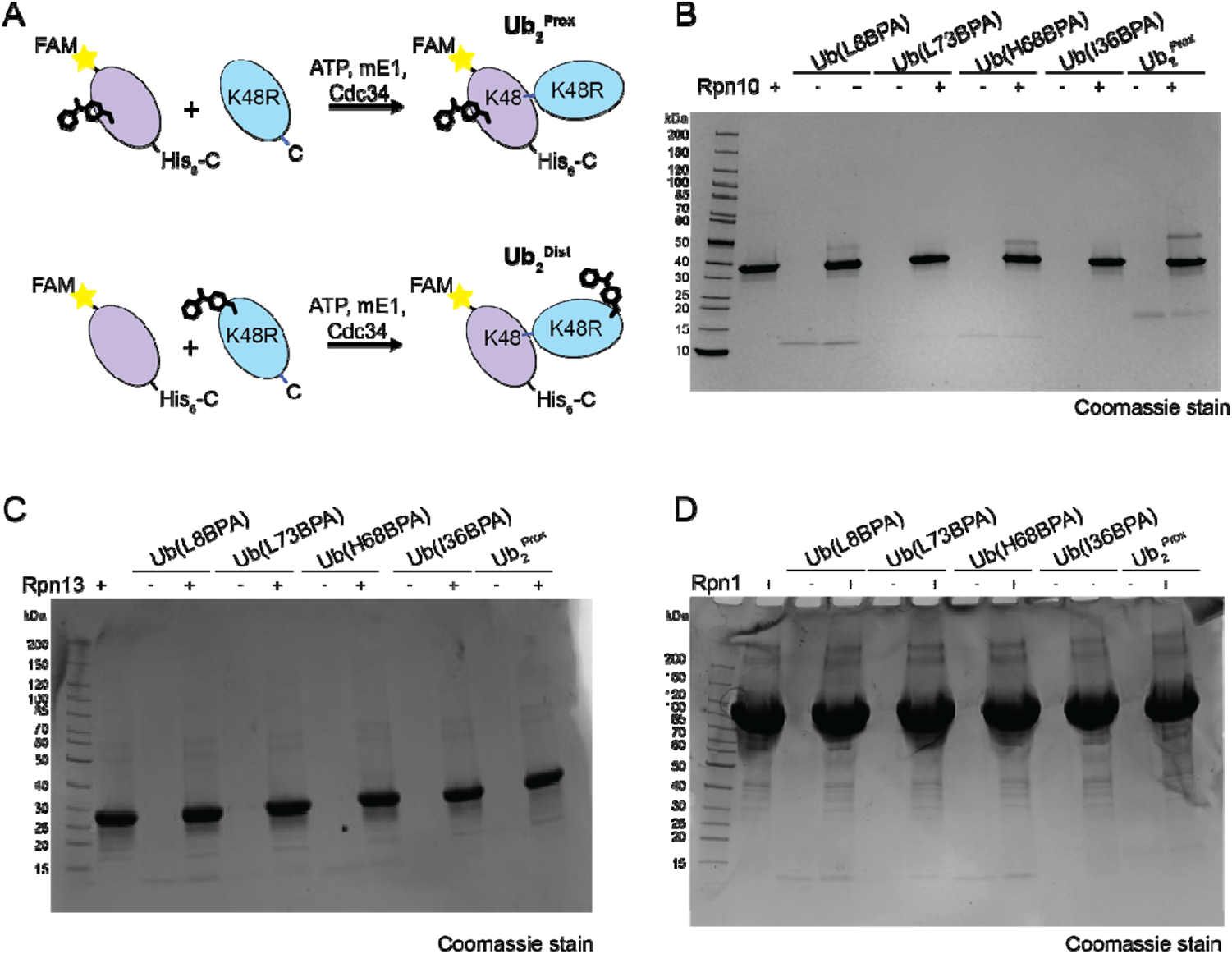
Generation and crosslinking of Ub(BPA) probes. **A)** Schematic of Ub_2_^Prox^ and Ub_2_^Dist^ probe production via Ub(BPA)-Ub conjugation, with Prox and Dist indicating the BPA position in the proximal or distal ubiquitin of the resulting K48-linked dimer. To direct the conjugation and prevent formation of larger oligomers, the proximal, FAM-labeled Ub carried a C-terminal His_6_ tag, while the distal Ub contained a K48R mutation. **B)** Coomassie-stained SDS-PAGE gel of Ub(L8BPA), Ub(L73BPA), Ub(H68BPA), Ub(I36BPA), and Ub_2_^Prox^ crosslinking to Rpn10, related to Figure 1F. **C)** Coomassie-stained SDS-PAGE gel of Ub(L8BPA), Ub(L73BPA), Ub(H68BPA), Ub(I36BPA), and Ub_2_^Prox^ crosslinking to Rpn13, related to Figure 1G. **D)** Coomassie-stained SDS-PAGE gel of Ub(L8BPA), Ub(L73BPA), Ub(H68BPA), Ub(I36BPA), and Ub_2_^Prox^ crosslinking to Rpn1, related to Figure 1H.

**Supplemental Figure 2:**
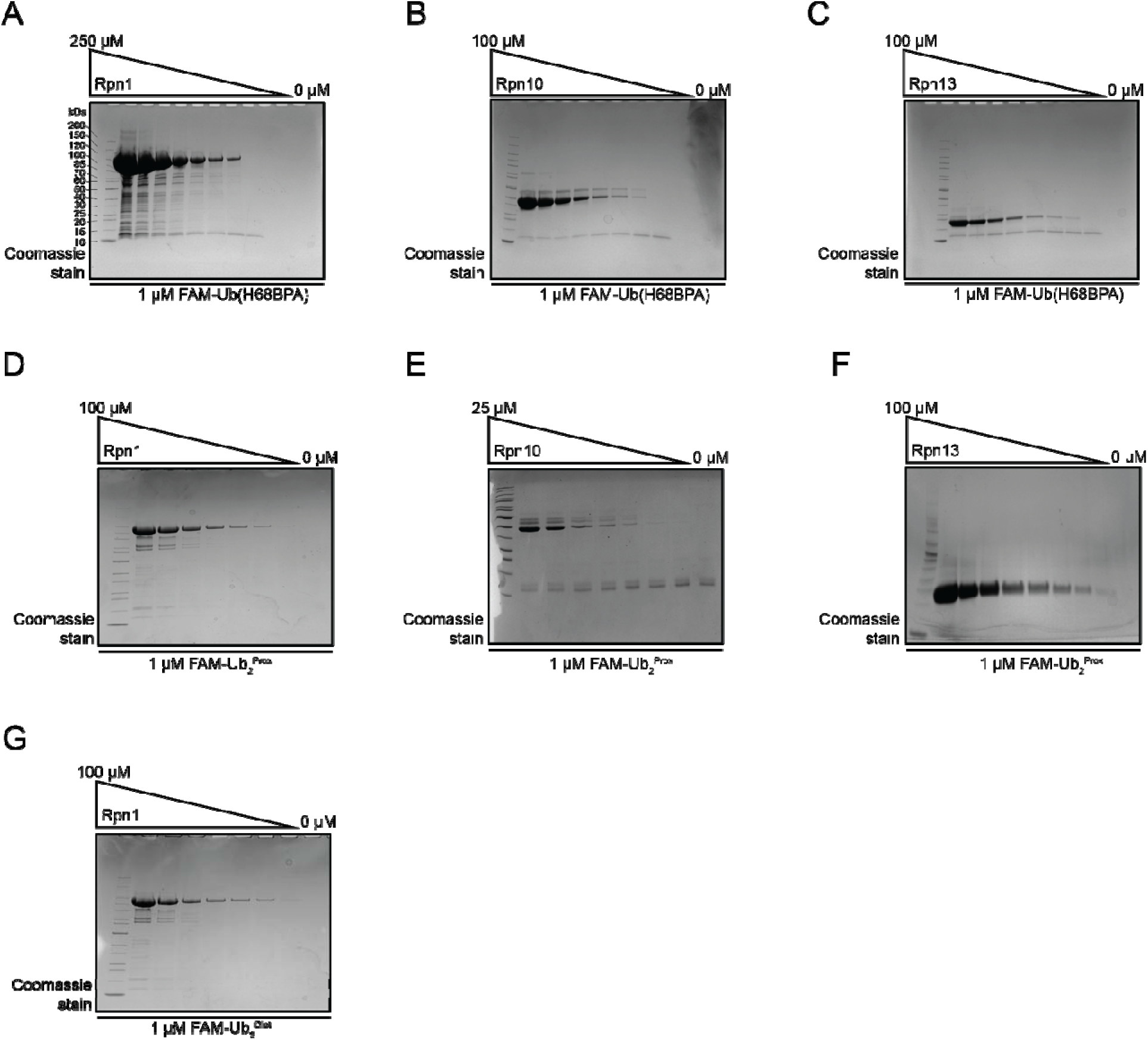
Coomassie-stained SDS-PAGE gels for the analyses of Ub(BPA) crosslinking to proteasomal receptors. **A)** Representative Coomassie-stained gel of Ub(H68BPA) crosslinked to various concentrations of Rpn1, related to Figure 2A. **B)** Representative Coomassie-stained gel of Ub(H68BPA) crosslinked to various concentrations of Rpn10, related to Figure 2B. **C)** Representative Coomassie-stained gel of Ub(H68BPA) crosslinked to various concentrations of Rpn13, related to Figure 2C. **D)** Representative Coomassie-stained gel of Ub_2_^Prox^ crosslinked to various concentrations of Rpn1, related to Figure 2D. **E)** Representative Coomassie-stained gel of Ub(H68BPA) crosslinked to various concentrations of Rpn10, related to Figure 2E. **F)** Representative Coomassie-stained gel of Ub(H68BPA) crosslinked to various concentrations of Rpn13, related to Figure 2F. **D)** Representative Coomassie-stained gel of Ub_2_^Dist^ crosslinked to various concentrations of Rpn1, related to Figure 2G.

**Supplemental Figure 3:**
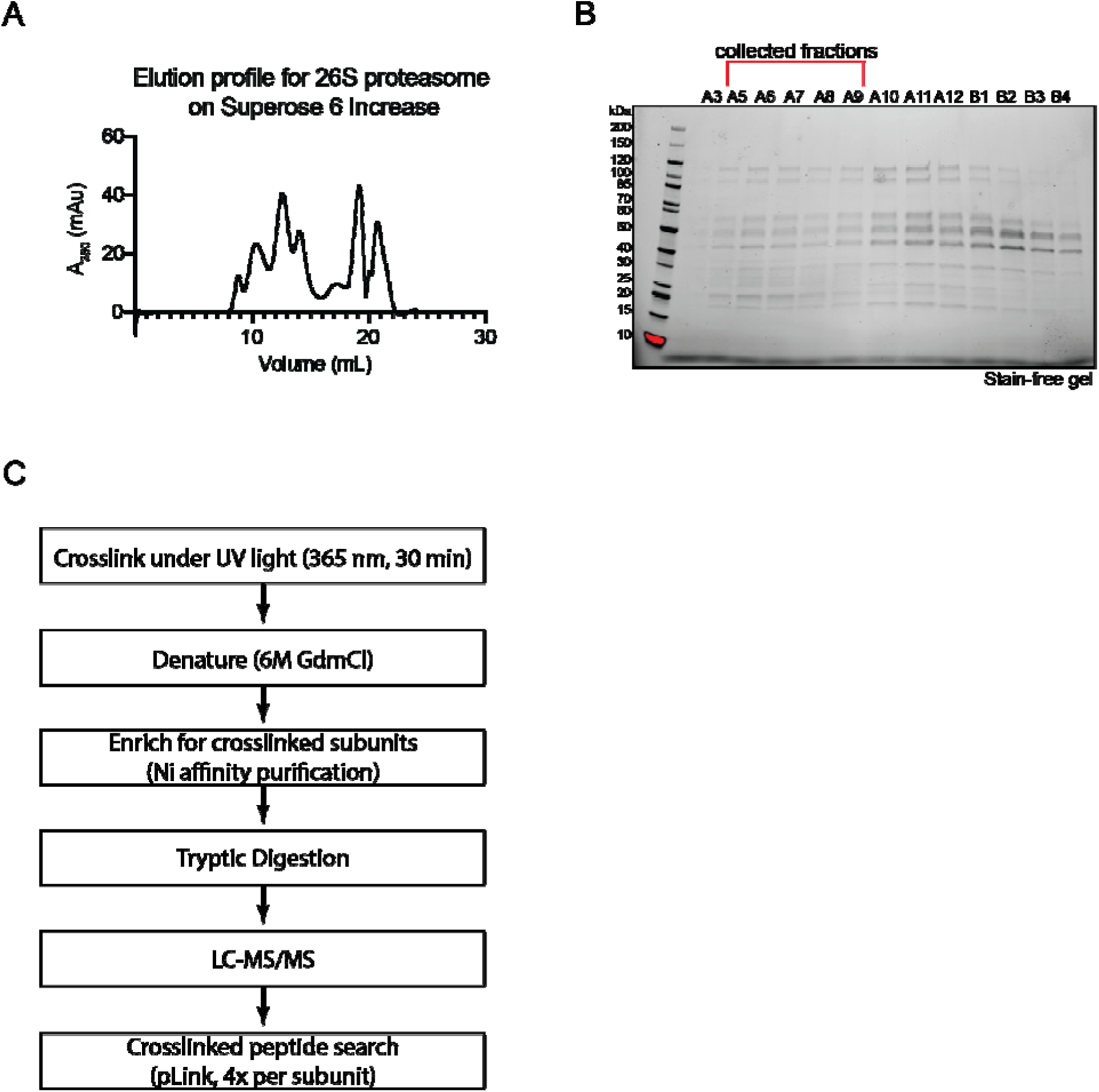
Purification and crosslinking of 26S proteasomes. **A)** Size-exclusion chromatogram of the yeast 26S proteasome eluting from a Superose 6 Increase column. **B)** Stain-free SDS-PAGE gel of fractions from size-exclusion chromatography purification of 26S proteasome shown in panel (A). Collected fractions are indicated above. **C)** Workflow for the photocrosslinking mass spectrometry experiments and identification of crosslinked peptides.

**Supplemental Figure 4:**
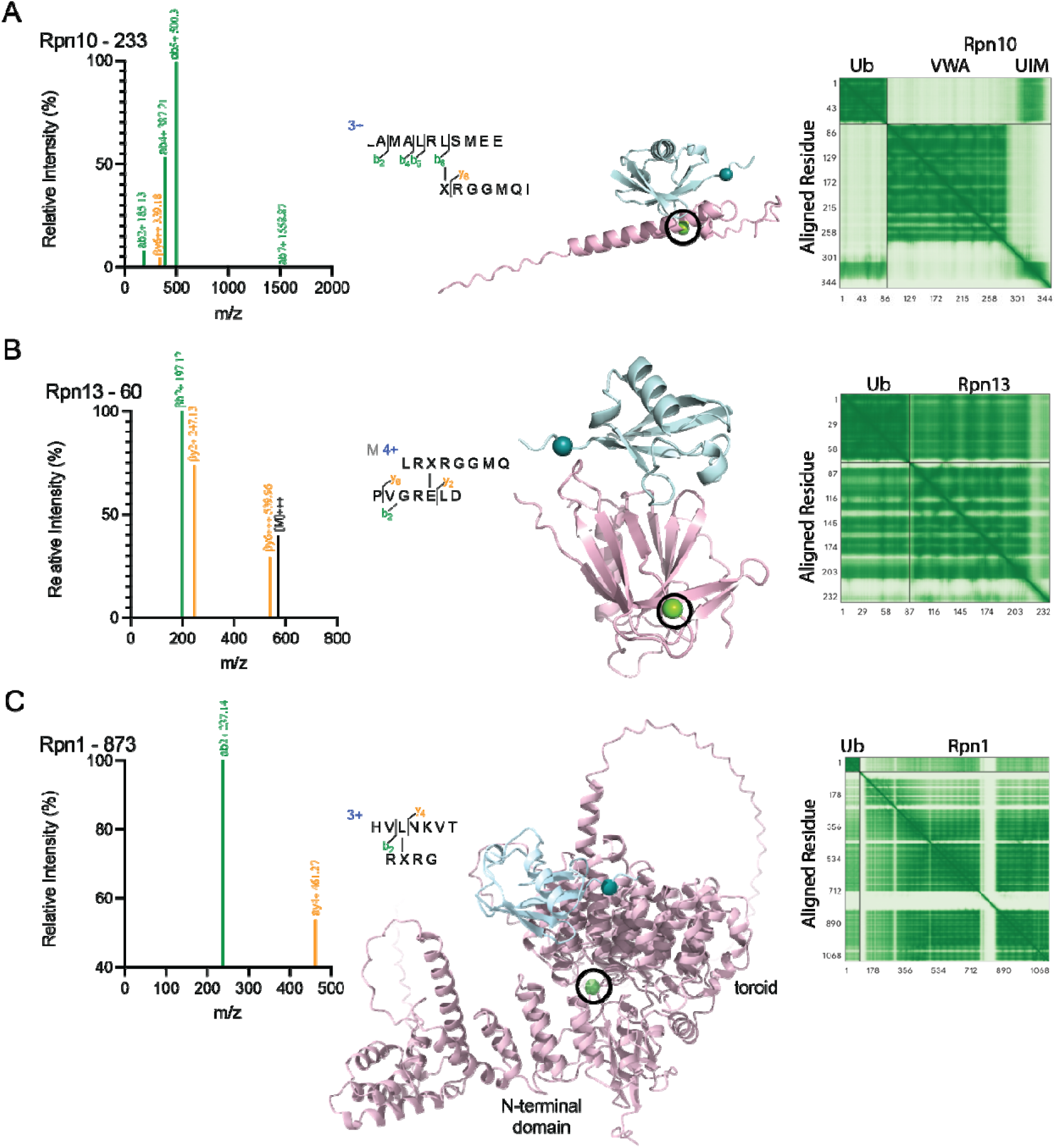
Ub(BPA)-receptor crosslink analyses. **A)** Mass spectrum of identified peptide containing a crosslink to position 233 in Rpn10 (left) and schematic of identified peptide (middle left). AlphaFold structural model of ubiquitin bound to Rpn10 (middle right), with crosslink site in Rpn10 shown as a green sphere and the BPA in Ub(L73BPA) indicated as a teal sphere. The AlphaFold predicted error plot is show on the right, with higher confidence residues colored green and lower confidence residues colored white. **B)** Mass spectrum of identified peptide containing a crosslink to position 60 of Rpn13 (left) and schematic of identified peptide (middle left). AlphaFold structural model of ubiquitin bound to Rpn13 (middle right), with crosslink site in Rpn13 shown as a green sphere and the BPA in Ub(L73BPA) indicated as a teal sphere. The AlphaFold predicted error plot is show on the right, with higher confidence residues colored green and lower confidence residues colored white. **C)** Mass spectrum of identified peptide containing a crosslink to position 873 of Rpn1 (left) and schematic of identified peptide (middle left). AlphaFold structural model of ubiquitin bound to the toroid domain of Rpn1 (middle right), with crosslink site L873 in a cleft between Rpn1’s toroid and N-terminal domain shown as a green sphere and the BPA position in Ub(L73BPA) shown as a teal sphere. The AlphaFold predicted error plot is shown on the right, with higher confidence residues colored green, and lower confidence residues colored white.

**Supplemental Figure 5:**
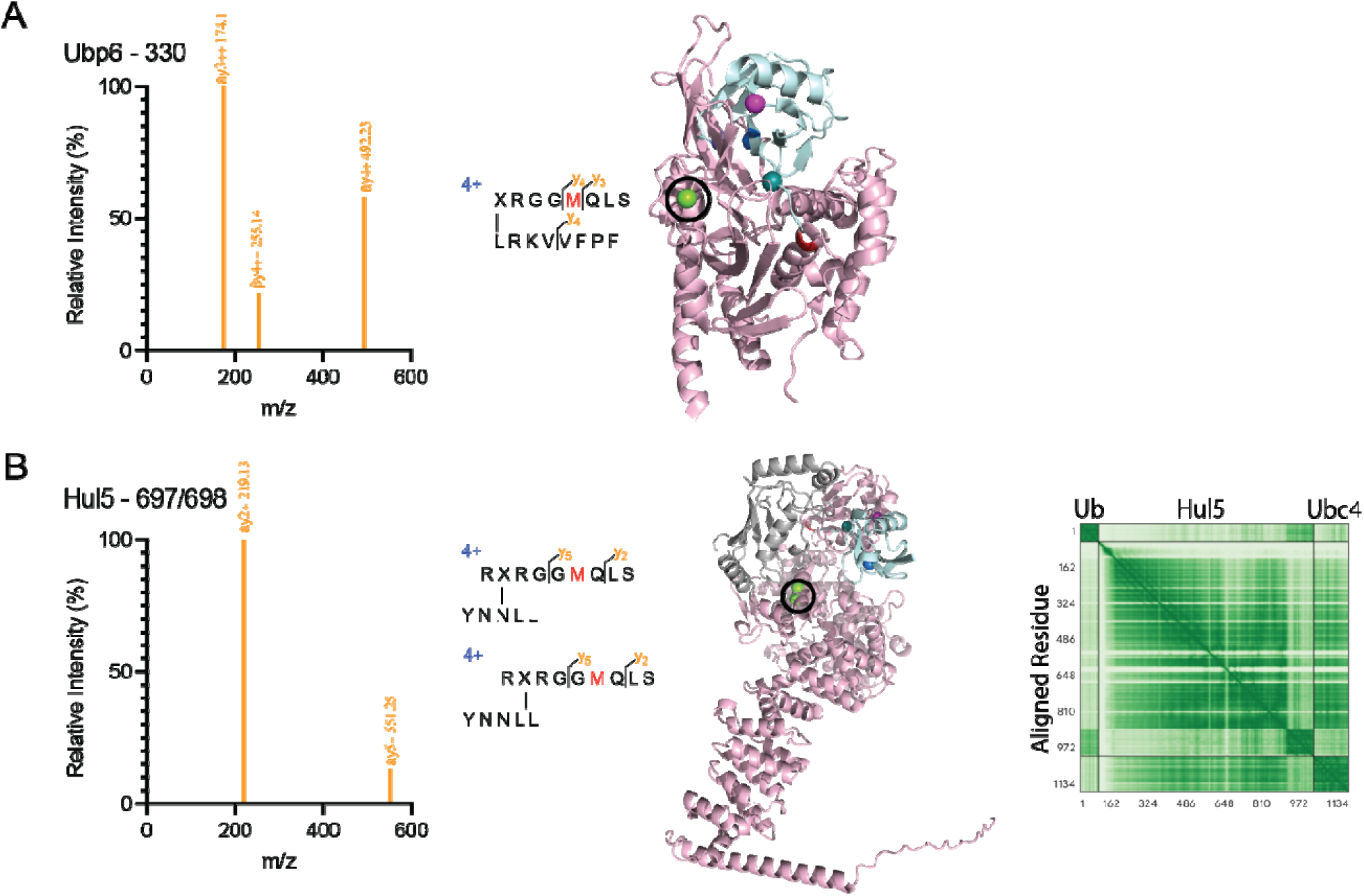
Crosslinks to proteasome-bound Ubp6 and Hul5. **A)** Mass spectrum of an identified peptide containing a crosslink at position 330 of Ubp6 (left) and schematic of identified peptide (middle). Structure of ubiquitin bound to Ubp6 (PDB: 7QO4), with crosslink site in Ubp6 shown as a green sphere and the BPA position in Ub(L73BPA) indicated as a teal sphere (right). **B)** Mass spectrum of identified peptide containing a crosslink to positions 697/698 of Hul5 (left) and schematic of identified peptide (middle left). AlphaFold structural model of ubiquitin bound to Hul5 (middle right), with crosslink site in Hul5 shown as a green sphere and the BPA in Ub(L73BPA) shown as a teal sphere. The AlphaFold predicted error plot is shown on the right, with higher confidence residues colored green and lower confidence residues colored white.

**Supplemental Figure 6:**
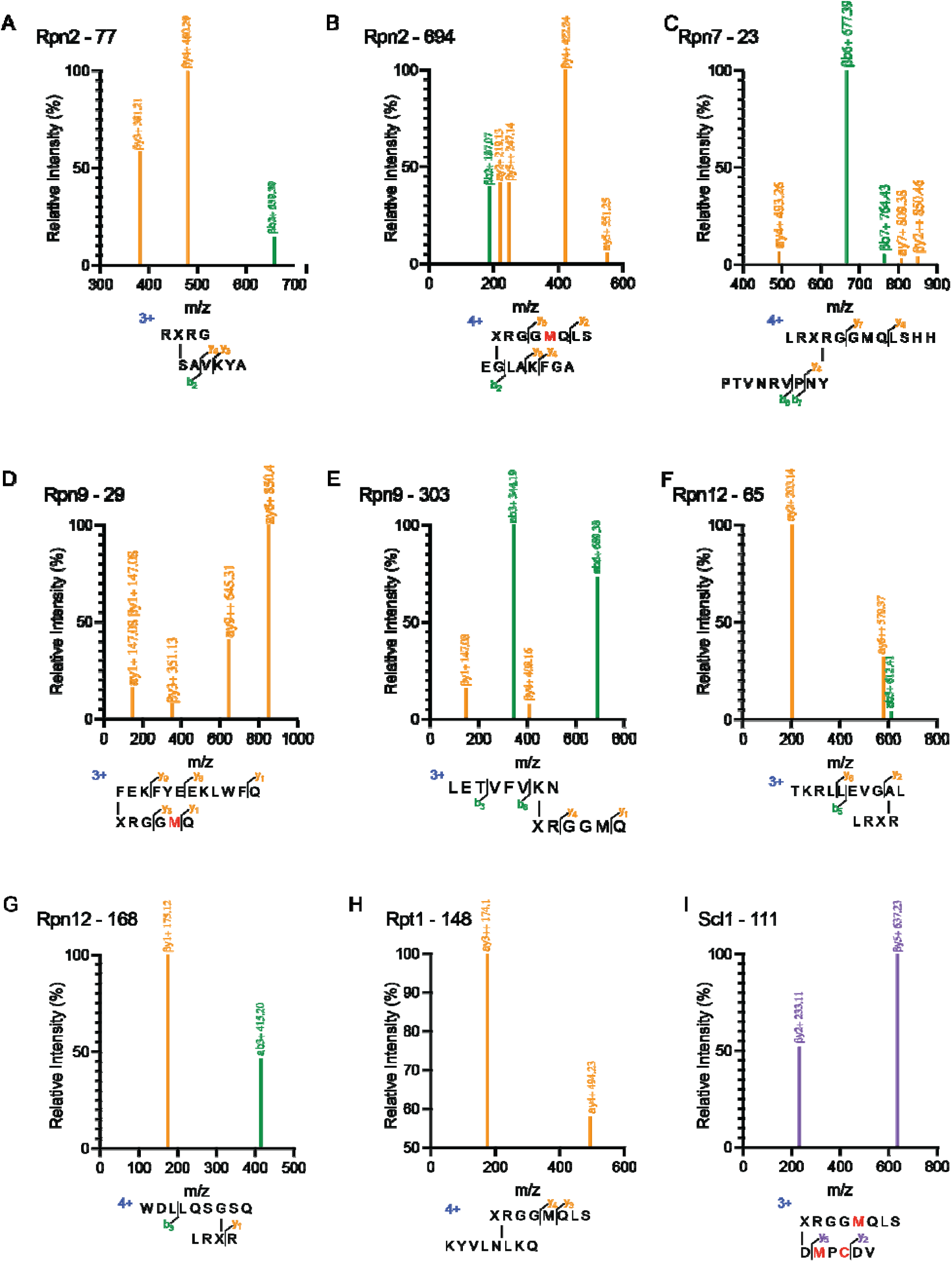
Mass spectra of crosslinked peptides from proteasomal subunits. **A)** Mass spectrum of an identified peptide containing a crosslink at position 77 in Rpn2 77 (top) and schematic of this peptide (bottom), with BPA indicated as ‘X’. **B)** Mass spectrum of an identified peptide containing a crosslink at position 294 in Rpn2 (top) and schematic of this peptide (bottom), with BPA indicated as ‘X’. **C)** Mass spectrum of an identified peptide containing a crosslink at position 23 in Rpn7 (top) and schematic of this peptide (bottom), with BPA indicated as ‘X’. **D)** Mass spectrum of an identified peptide containing a crosslink at position 29 in Rpn9 (top) and schematic of this peptide (bottom), with BPA indicated as ‘X’. **E)** Mass spectrum of an identified peptide containing a crosslink at position 303 of Rpn9 (top) and schematic of this peptide (bottom), with BPA indicated as ‘X’. **F)** Mass spectrum of an identified peptide containing a crosslink at position 65 of Rpn12 (top) and schematic of this peptide (bottom), with BPA indicated as ‘X’. **G)** Mass spectrum of an identified peptide containing a crosslink at position 168 of Rpn12 (top) and schematic of this peptide (bottom), with BPA indicated as ‘X’. **H)** Mass spectrum of an identified peptide containing a crosslink at position 148 of Rpt1 (top) and schematic of this peptide (bottom), with BPA indicated as ‘X’. **I)** Mass spectrum of an identified peptide containing a crosslink at position 111 of Scl1 (top) and schematic of identified peptide (bottom), with BPA indicated as ‘X’.

